# A single-cell resolved cell-cell communication model explains lineage commitment in hematopoiesis

**DOI:** 10.1101/2021.03.31.437948

**Authors:** Megan K. Franke, Adam L. MacLean

## Abstract

Cells do not function in isolation. Arguably, every cell fate decision occurs in response to environmental signals. In many cases cell-cell communication alters the dynamics of a cell’s internal gene regulatory network to initiate cell fate transitions, yet models rarely take this into account. Here we develop a multiscale perspective to study the granulocyte-monocyte vs. megakaryocyte-erythrocyte fate decisions. This transition is dictated by the GATA1-PU.1 network, a classical example of a bistable cell fate system. We show that, for a wide range of cell communication topologies, even subtle changes in signaling can have pronounced effects on cell fate decisions. We go on to show how cell-cell coupling through signaling can spontaneously break the symmetry of a homogenous cell population. Noise, both intrinsic and extrinsic, shapes the decision landscape profoundly, and affects the transcriptional dynamics underlying this important hematopoietic cell fate decision-making system.

## Introduction

The production of mature cell types from stem cells and progenitors is essential for development and organ homeostasis. Nevertheless, in few cases are we able to fully specify the conditions necessary to drive stem cell differentiation towards a particular cell lineage. Stem cell differentiation is controlled by cell-internal and external signals that, in turn, control the transcriptional state of the cell and specify its eventual fate [1–3]. These changes in transcriptional state are often irreversible and involve binary choices. Thus, multipotent cells, through successive binary lineage specification choices, eventually become committed to a specific lineage and cell fate. Significant unanswered questions remain regarding the cell fate decisions that dictate lineage specification of stem/progenitor cells: how large is the role of extracellular signaling in regulating cell differentiation? What mechanisms allow cells from initially homogeneous or clonal populations to converge to different lineages? How does one population of progenitor cells maintain stable heterogeneous subpopulations of committed cell types?

During lineage specification, changes in gene expression are controlled by gene regulatory networks (GRNs), consisting of genes and their protein products (transcription factors), which are able to regulate the expression of various genes, including other transcription factors, creating feedback loops [4]. Codified through mathematical models, GRNs can be studied in light of their steady states, where each steady state can represent a committed cell fate. Certain GRN topologies permit bistability, that is, more than one steady state can be reached for a single set of biological conditions [5]: such topologies are frequently observed in networks that control cell fate decisions [6, 7]. The GRN topology that dictates a particular lineage decision is generally conserved across cells, thereby providing insight into the intracellular dynamics that occur during cell differentiation. Crucially, the GRNs that instigate or reinforce lineage decisions are not only controlled by cell-intrinsic networks, but also by cell-extrinsic signals.

The GRN topology considered in this work consists of two mutually repressive genes; a topology frequently observed among gene networks mediating lineage decisions [8, 9]. One such lineage decision occurs during hematopoiesis: myeloid progenitor cells make a binary choice between commitment to the megakaryocyte-erythroid (ME) lineage or the granulocyte-monocyte (GM) lineage. *GATA1* and *PU.1* (*SPI1*) mutually inhibit one another; *GATA1* is expressed in the ME lineage and *PU.1* is expressed in the GM lineage. This mutually repressive GRN has been extensively studied and characterized in models, mostly consisting of ordinary differential equations that permit bistability, thus enabling investigation into the dynamics of this myeloid lineage decision [10–14]

Given the *GATA1-PU.1* mutual inhibitory loop that leads to bistability, changing the initial conditions (gene expression levels) within a bistable region is sufficient to change the cell fate. It has thus been proposed that random fluctuations of *GATA1* and *PU.1* levels are primarily responsible for determining cell fate in the bipotent progenitor population that has dual ME and GM lineage potential [13]. More recently this notion was challenged by a study that used a double reporter mouse (PU.1^eGFP^Gata1^mCherry^) to show that that random fluctuations of *PU.1* and *GATA1* are insufficient to initiate the cell fate decision between ME and GM lineages [15]. Hoppe et al. also provide strong evidence that ME v. GM lineage specification cannot be determined solely from initial ratios of *PU.1* to *GATA1* expression.

As noted by Hoppe et al. [15], the role of extracellular signaling could resolve this controversy. However, the role of extracellular signaling has been chronically understudied in models of gene regulatory networks and their stability. Extracellular signaling through paracrine factors enables populations of cells to share information and coordinate behaviors over short distances, and is implicated in a range of cell fate behaviors from developmental branching [16, 17] to patterning [18] and migration [19]. We begin this study with a simple question: how does short distance intercellular signaling impact myeloid lineage specification controlled by the *GATA1-PU.1* gene regulatory network?

Several studies have characterized the impact on cell fate of cell-cell communication through extra-cellular signals. One approach is to model the internal gene dynamics with differential equations and model cell-cell communication by allowing for molecular diffusion of proteins between neighboring cells [20–22]. These methods however must neglect anisotropic effects, and assume that signaling between cells occurs on the same timescale as intracellular dynamics: these processes can occur on vastly different timescales. Other approaches have relaxed this assumption, permitting different timescales by modeling the response time of a cell to a signal received as a random variable [23]. This approach however omits any detail of the intracellular dynamics, which are often crucial to the response; GRNs are the enforcing mechanisms of the phenotypic switches. Therefore, in order to model the behaviors of a population of communicating cells, the gap between intracellular dynamics and extracellular signaling must be bridged. A model of cell fate decision-making is needed that permits cell-cell communication while allowing for both description of the cell-internal dynamics and of the extracellular signaling dynamics without making diffusion-like approximations.

We present here a multiscale model that bridges this gap. We will assume deterministic dynamics within each cell and thus model cell-internal GRNs with ordinary differential equations (ODEs). We assume that signals sent between cells can be described by a Poisson process. Signals received by cells can then alter the internal GRN dynamics by changing ODE parameters. We test this model in a large range of different intercellular signaling topologies. We find that the addition of cell-cell communication to GRN dynamics leads to model outcomes (cell fate distributions) becoming probabilistic, with cell fate choice probabilities that depend on the cell’s position in a particular signaling topology. We discover that the model can intuitively characterize cell-cell coupling, changes to which impact the cell fate decisions made, which can lead to mixtures of heterogeneous cell types within a population. Finally, we study how noise propagates across various signaling topologies, and show that although both intrinsic and extrinsic noise alter the cell fate decision-making boundaries, extrinsic noise is the dominant driver of cell fate variability.

## Methods

### A multiscale model of cell-cell communication between single cells

To investigate how external signaling impacts intracellular dynamics during cell fate decisions, we first must select an internal GRN topology to model. We choose to model a mutual inhibitory GRN: such network topologies arise frequently in cell fate decision making and developmental biology [24, 25]. Within certain parameter regions, such models give rise to two stable steady states and bistability (Fig. 1A). For example, it has been observed that the GRN determining the bifurcation of erythroid/megakaryocyte lineages from granulocyte/monocyte lineages contains a core mutual inhibitory loop topology consisting of mutual antagonism of *GATA1* (*G*) and *PU.1* (*P*). For this genetic switch, high expression of *G* corresponds to commitment to the erythroid/megakaryocyte lineage and high expression of *P* corresponds to commitment to the granulocyte/monocyte lineage. This myeloid progenitor cell lineage decision has been rigorously studied, and there exist robust models of the intracellular GRN dynamics [10, 11, 13, 14, 26, 27]. Here we implement the ODE model defined by Chickarmane et al., given by Eqs. 1–3 [10]. These ODEs give rise to two stable steady states for a region of parameter space with respect to the parameters *A, B* and *C*, where *A, B*, and *C* are parameters summarizing environmental signals (Fig. 1B). The parameters through which the extracellular environment is summarized will be modified below to implement a multiscale model of cell-cell communication. The parameter values used throughout this work can be found in Table S1.

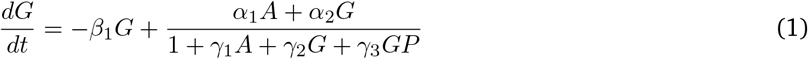

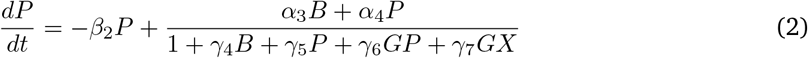

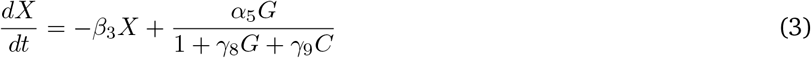

**Figure 1:**
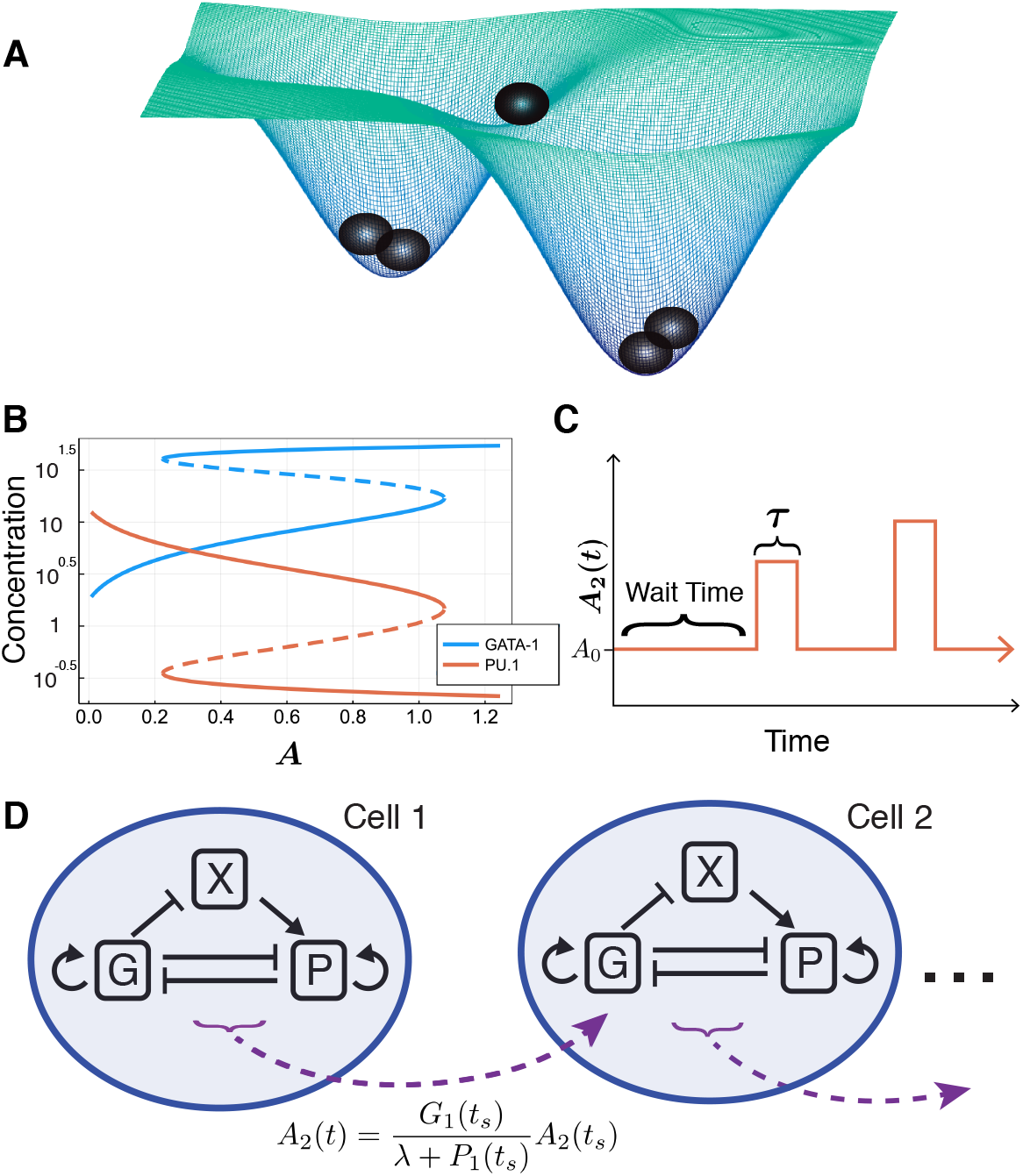
**(A)** Illustration of cells in a two-attractor state model (such as the GATA1-PU.1 inhibitory loop). **(B)** Bifurcation diagram for the system of ODEs described by Eqs. 1–3, with respect to the external signaling parameter *A*. **(C)** Cell-cell communication via signals sent between cells (e.g. from cell 1 to cell 2) is modeled by a Poisson process; with wait times sampled from an exponential distribution and fixed signaling ‘pulses’ of length *τ*, where the value of *A*_2_ is set by Eqs. 4 or 5. **(D)** Full model schematic, depicting the cell-internal gene regulatory network of {G,P,X} modeled by ODEs and the external signal *A*_2_ modeled by the Poisson process in (C).

To incorporate cell-cell signaling into the model, we must modify the cell-internal ODEs in a way that reflects the signal received by the cell without unrealistically changing the internal dynamics (e.g. a loss of bistability). For simplicity, we choose a single parameter *A* to summarize the effects of external signaling. We study both signals that recruit nearby cells to commit to the same lineage and the opposite lineage of the sender. For the first case, we consider two cells, cell 1 and cell 2, where cell 1 signals to cell 2. Let *G*_1_(*t*) and *P*_1_(*t*) denote the concentrations of *G* and *P* in cell 1 at time *t* respectively and let *A*_2_(*t*) be the value of the parameter *A* in cell 2 at time *t*. Since cell 1 signals to cell 2 to coordinate fate decisions (“be like me” signal), then *A*_2_(*t*) should increase if the ratio *G*_1_(*t*) : *P*_1_(*t*) indicates that cell 1 has committed to – or is increasingly likely to commit to – the ME state (where *G* is highly expressed). Conversely, *A*_2_(*t*) should decrease if the ratio *G*_1_(*t*) : *P*_1_(*t*) indicates that cell 1 has committed to – or is increasingly likely to commit to – the GM state where *P* is highly expressed. Based on these assumptions, we constructed a communication model: we treat signaling as a Poisson process, where the wait time between signals being sent by a cell follows an exponential random variable with mean *μ*. After this wait time, at time *t_s_*, a signal is sent and we change the value of *A*_2_ by:

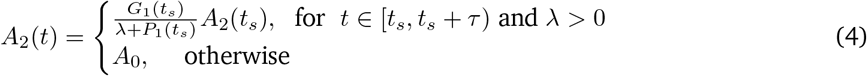

After a delay *τ*, *A_2_* returns to it’s original value, *A*_0_ (Fig. 1C). Returning to the same value *A*_0_ between signals ensures that attractor states (Fig. 1B) do not change. In Fig. 1D is a schematic of this signaling model. Fig. S1 gives a sample simulation of a chain of four cells and the values of *A*_2_, *A*_3_, and *A*_4_ plotted over time. In the case of signals that promote nearby cells to commit to the opposite lineage, we can follow the same principles to define a signal:

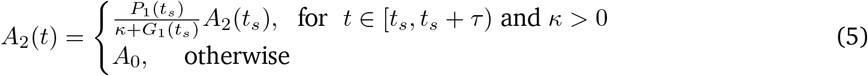

These definitions of signaling also allow for multiple cells to signal a single cell at similar times while still having *A*_2_ well defined. We implemented this model in Julia [28], where we used the DifferentialEquations.jl [29] and Distributions.jl [30] packages for numerical simulation.

The model developed here shares similarities with models specified by piecewise-deterministic Markov processes (PDMPs), first introduced by Davis [31]. Such models are comprised of continuous dynamics and a discrete Markov process which can alter the continuous dynamics at discrete time points. PDMPs have previously been applied to biological systems for the study of gene expression and genetic networks [32, 33]. In the case of two cells sharing one directed signal between them, the model described here closely resembles a PDMP, with the exception that here, due to information transfer between cells, the discrete process is not Markovian.

### Model specification and choice of parameters

To elucidate our choice of signaling model, we briefly discuss some alternatives. Examples of alternative signaling model choices include: signals that change more than one ODE parameter, additive rather than multiplicative signaling, or signals that permanently change ODE parameters. We choose to modify a single parameter for interpretability and to constrain the model space: the ODE system exhibits bifurcations with respect to many of its parameters, and we do not seek to explore bifurcations of codimension 2 or greater here. Modeling with additive signals brings complications, such as the possibility of obtaining negative values of *A*. Also, if a large number of committed cells are communicating with a single cell, the value of |*A*| may be unrealistically large and dominate the ODE. This problem is dealt with in the signaling model we present here through the parameter λ. Lastly, permitting signals to cause permanent changes to ODE parameters (rather than over time intervals) can result in the divergence of *A*. Thus, we have selected a signaling model that can capture how changes in the extracellular environment can alter cell internal GRN dynamics while still preserving the overall behaviors (i.e. the attractor states) of the dynamical system.

In this work, we will assume that cells are homogeneous, i.e. all cells share the initial conditions (*G, P, X*) = (20, 1, 20). The internal GRNs are identical with the same parameter values, including the same initial value of *A*, denoted *A*_0_. We set the signaling period to be *τ* = 5, and the mean of the signaling wait time distribution to be *μ* = 50. Changing the values of *μ* and *τ* does cause qualitative changes as long as *τ* is sufficiently large relative to *μ*. For more insight on the relationship between the wait time distribution and *τ*, see Fig. S2. Unless otherwise specified, we set λ = *κ* = 1. We initially chose λ = *κ* = 1 to avoid unrealistic behaviors when *G* or *P* ≈ 0. We will provide an in depth discussion on the roles of these parameters in the next section. For all topologies depicted schematically below, regular arrowhead arrows correspond to consensus signaling (Eq. 4) and flat (inhibition) arrowhead arrows correspond to dissensus signaling (Eq. 5).

As a result of signaling, cell fate decisions became probabilistic rather than deterministic. Then, for each cell in a signaling topology, we can examine the probability distribution of the cell converging to a certain lineage. To do so, for each signaling topology tested, we selected a range of values of *A*_0_ to simulate. The step size between the *A*_0_ values was scaled between each topology depending on the range of the probability distribution. For each value of *A*_0_, we simulated *N* trajectories, counting the number of times each cell converged to both fates. From these counts, we estimated the probability that each cell converged to the *G* high state and plotted:

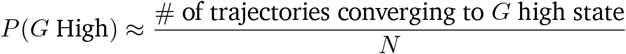

*A* sample of the simulated data points can be found in Fig. S3. Unless stated otherwise, probabilities are estimated by running *N* = 1, 000 simulations.

While λ = 1 was an intuitive first choice to avoid divergence in signaling parameters, it relies on the assumption that a cell converging to the *P* high state implies that *P* is more highly expressed than *G* + 1 and vice versa. This turns out to not necessarily be true. Rather, for a given value of *A*_0_, being in the *P* high state means that *P* is more highly expressed relative to the other stable steady state value of *P*. Recalling Fig. 1B, we see that for some values of *A*_0_, for example *A*_0_ = 0.8, *G* is always more highly expressed than *P* regardless of which stable steady state a cell converges to. Looking at the bifurcation diagram, we see that in the *P* high state, the maximum steady state value of *G* is approximately 16.97 and the minimum value of *P* is approximately 1.475. Similarly, in the *G* high state, the maximum steady state value of *P* is approximately 0.3529 and the minimum value of *G* is approximately 40.35. From here, we wanted to select values of λ which satisfy the following:

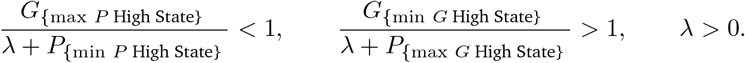

These inequalities are satisfied for λ ∈ [15.495, 39.997]. We will explore how selecting values of λ in or near this range changes the behaviors of both individual cells and populations of cells.

### Intrinsic and extrinsic cell-cell communication noise

In this work, we focus on how intercellular signaling alters cell decision making. Therefore, we only consider noise with respect to signaling (not the internal GRN dynamics). Noise in the signal can arise due to the noisy extracellular environment (extrinsic noise) or can arise during the signal transduction within a cell (intrinsic noise) [22, 23]. Here we define models for both sources of noise.

For extrinsic noise, i.e. noise with respect to the extracellular environment, let *η_e_* be a random variable such that 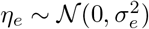, where 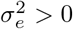. For Cell *k* in a given topology, at the start of each new wait period, we sample a value of *η_e_* and update *A_k_*(*t*) by

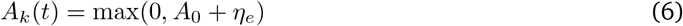

We truncate *A_k_*(*t*) at zero to avoid negative values. However, we select values of 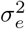 small enough that the truncation is rarely necessary in simulation. Note that noise in *A_k_*(*t*) results in noise during the signaling period as well by Eqs. (4) and (5).

For intrinsic noise, i.e. noise with respect to signal transduction, let *η_p_* be a random variable such that 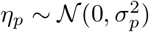, where 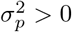. For Cell *k* in a given topology, at the start of each new pulse period, we sample a new value of *η* and update *A_k_*(*t*).

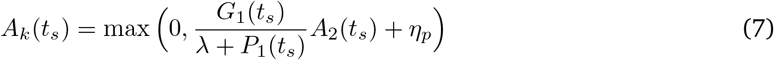

In the analysis of noise effects below, we decrease the mean wait time to *μ* = 40, in order to distinguish between the effects of intrinsic, extrinsic, or no added noise.

## Results

### Cell-cell communication over a diverse range of signaling topologies leads to divergent cell fates

We begin by testing the base model behavior for different signaling topologies. We sought to ensure that core assumptions were upheld, namely that all cells eventually reach a steady state, even in the presence of noisy signaling, and that cells exhibit bistability with respect to *A*_0_ (the initial value of *A*). These behaviors were preserved in all signaling topologies tested. Moreover, we found that the addition of cell-cell signaling to a previously deterministic model results in probabilistic cell fate choices.

In Fig. 2, simulated trajectories for a 20-cell loop topology are shown. We plot only the concentration of *GATA1* (G) as a proxy for fate choice, since its state is sufficient to determine the steady state of the full system (if *G* is high at steady state, *P* is low, and vice versa); as illustrated by the trajectories of both *G* and *P* (Fig. S4). These trajectories illustrate the variety of cell fate distributions that are observed in the presence of cell-cell communication (fate convergence to either the high or low state, and fate divergence). In Fig. 2C-D, the values of *A*_0_ are equal, demonstrating that *A*_0_ does not determine the proportion of cells in each state. Rather, for each value of *A*_0_, the probability of each cell converging to a certain state can be computed.

**Figure 2:**
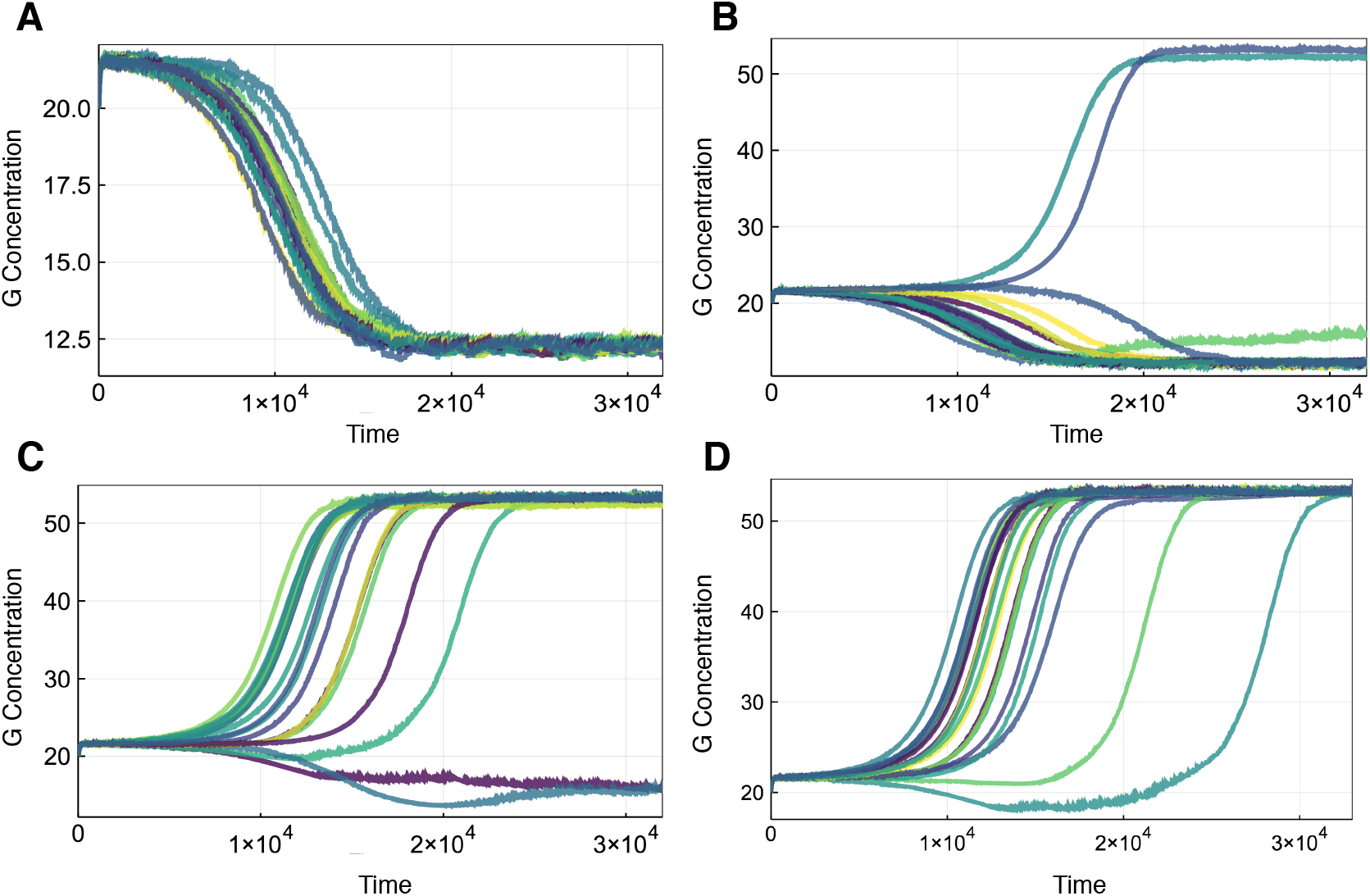
Sample trajectories of the concentration of species *G* over time for each cell in a loop of 20 cells with λ = 28. Colors used only to distinguish the trajectories. **(A)** *A*_0_ = 0.995. **(C-D)** *A*_0_ = 1.0.

Investigating how signals propagate down a chain of cells, two striking observations are made. First, each additional signal shifts the *P*(*G*) curve (the probability of reaching the *G* high state) to the left, although these shifts are successively smaller down the chain such that eventually the distributions converge. Second, a “ sharpening” of the *P*(*G*) distributions occurs whereby the region of uncertain fate decreases for cells further down a chain (Fig. 3A). This is a hallmark of cooperativity: accumulated signals by cell-cell communication act to reinforce cell fate decision-making, leading to regions of uncertainty that are decreasing with the total number of signals that are acting on a cell.

**Figure 3:**
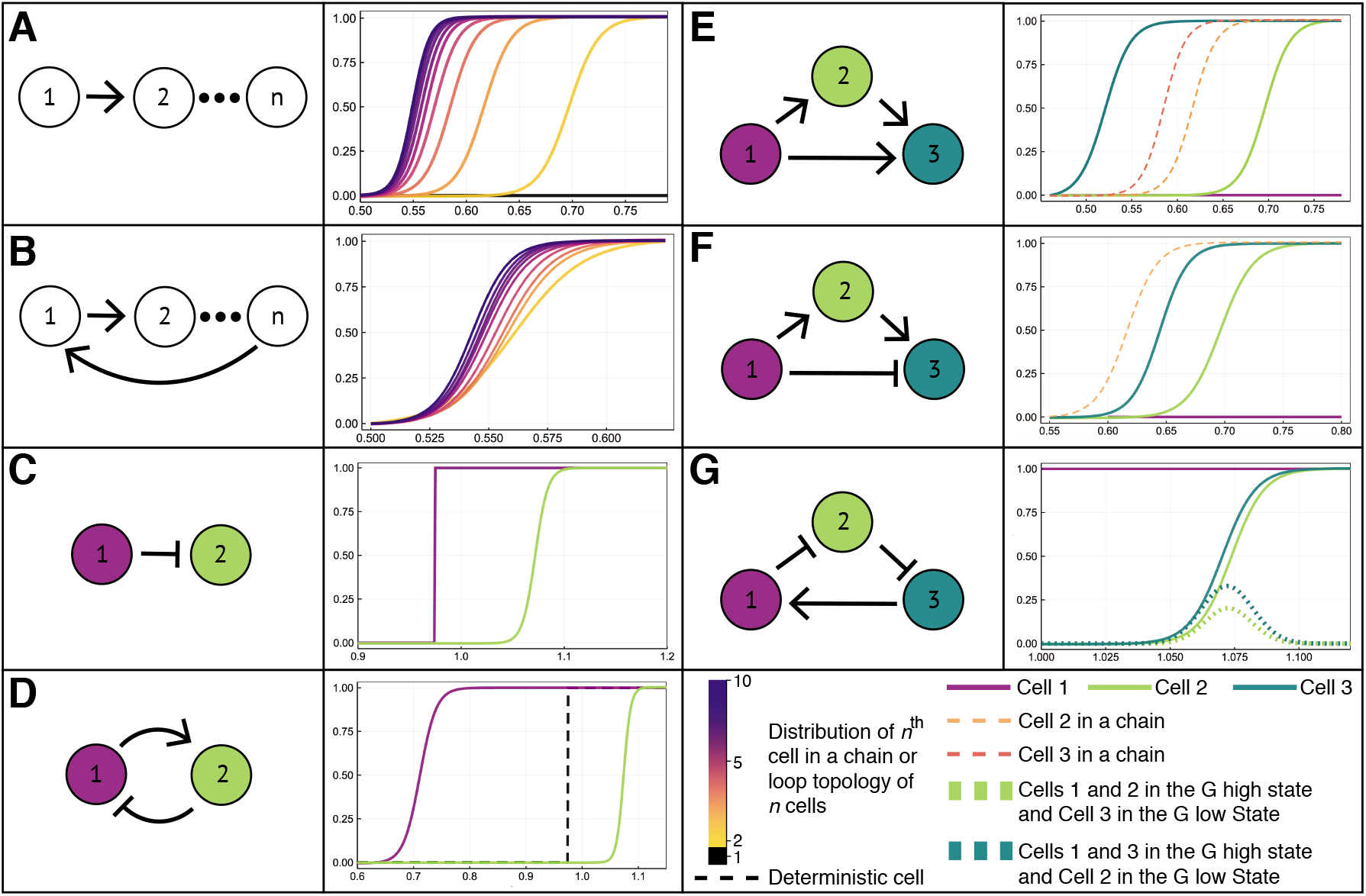
Schematic of different cell signaling topologies (left) and the corresponding probability distributions: the probability that each cell in the network will commit to the *G* high steady state for different values of *A*_0_. **(A)** Chains of cells of length *n*. Each probability distribution corresponds to one cell in the chain. **(B)** Loops of cells of size *n*. In a loop of fixed size, each cell from 2 to *n* has the same probability distribution; thus one curve is plotted for each loop of *n* cells. **(C)** A chain of two cells with an dissensus signal: the distribution of cell 2 fates is shifted to the right, i.e. it commits at higher values, compared with consensus signal in (A) where the distribution of cell 2 is shifted to the left. **(D)** A loop of two cells with one consensus and one dissensus signal; dashed line denotes the distribution of a single cell that receives no signals (deterministic). Note that we cannot observe the fate where cell 1 is in the *G* low state and cell 2 is in the *G* high state. **(E)** Three-cell feed forward motif; cells 3 (dashed orange) and 4 (dashed red) from a chain of cells marked for comparison. We see that receiving multiple signals results in a non-additive synergistic effect. **(F)** Feed forward motif with a dissensus signal; cell 3 (dashed orange) from a chain of cells marked for comparison. **(G)** Loop of three cells with two dissensus signals. Dashed lines denote the probabilities that: i) cells 1 & 2 are in *G* high state while cell 3 is in *G* low state (dark green); or ii) cells 1 and 3 are in *G* high state while cell 2 is in the *G* low state (light green).

In loops, all cells converge to the same fate for each simulated trajectory, therefore every cell in the model has the same probability distribution. (Below we will see that the coordination of fates in all cells within a loop signaling topology can be broken by changing the cell coupling strength.) The probability distribution for a cell loop depends on the number of cells in the loop, and we see that for loops of size *n*, both of the behaviors observed above are recapitulated (Fig. 3B). Moreover, the limiting distributions for the cell chain and the cell loop are the same, highlighting an important underlying convergence.

If we consider a dissensus signal (the opposite of a consensus signal), the fate probability distribution of cell 2 shifts in the opposite direction than previously (Fig. 3C), relative to the value of *A*_0_ at which cell 1 switches lineages. When an inhibition signal is incorporated into a loop (Fig. 3D), the two cells in the loop no longer coordinate lineages for every trajectory as they did when all signals were regular. Notably, here there exists a large range of *A*_0_ values where cell 1 converges to the *G* high state and cell 2 converges to the *G* low state with probability one. Thus, the inhibition signals significantly changes the resulting behaviors in both chains and loops of cells.

For signaling topologies consisting of three cells, concordant effects due to cell-cell communication are found. In a feed forward signaling motif (Fig. 3E), multiple promoting signals reinforce the lineage choice of cell 3, thus increasing cooperativity and as a result the cell commits to the *G* high state at smaller values of *A*_0_ than it would in a cell chain. Other three-cell signaling topologies also display interesting behaviors (Fig. 3F-G and Fig. S5). For an inhibitory feed forward motif, we observe a multiplicative effect between the inhibitory signal from cell 1 and the activating signal received indirectly from cell 1 via cell 2 (Fig. 3F). The addition of the inhibition signal results in the distribution of cell 3 being shifted to the right relative to a corresponding cell in a chain. For a double inhibitory topology (Fig. 3G), we find that for a region of *A*_0_ the cell fates can diverge (two cells in the high state; the third in the low state) with nonzero probability.

While these results pertain to small and idealized cell signaling networks, they all point to a general and important conclusion. In each of the above experiments, the addition of cell-cell signaling transformed a population of homogeneous, independent, and deterministic cells into one of heterogeneous cells that choose fates non-deterministically. This shows that, given cells with identical initial conditions (i.e. transcriptional states), external signaling changes cell fate outcomes at the population level. This corroborates a major finding of Hoppe et al. [15]: that GMP/MEP cell fate decisions are not predictable from initial transcription factor ratios alone. Our model not only supports this result, but offers an explanation. The missing determinant of cell fate commitment – which acts to break the symmetry of an initially homogenous population of cells – is cell-cell communication.

### Varying the strength of cell-cell coupling results in different ratios of stable subpopulations of heterogeneous cell types

Through deeper investigation of cell-cell communication in the model framework introduced above, we found that divergence of cell fates can occur under the control of the model parameter λ. We thus describe λ as a “cell-cell decoupling” mechanism. Analysis of cell-cell coupling and decoupling led to two significant findings: cell-cell communication can explain how a population of cells maintains stable subpopulations of heterogeneous cell types; and the distribution of cells that converge to each fate depends both on cell-cell coupling and on the external environment.

First, we assessed how different values of λ change the simplest signaling topology: a chain of two cells (Fig. 4A). As λ increases, the probability distribution of the second cell shifts to the right. Note that λ = 22 is close to a critical value of λ, resulting in near-deterministic cell fate decision-making. The value of *A*_0_ where the curve for λ = 22 switches from one lineage to another is extremely close to the value at which a deterministic cell with the same initial conditions switches. Overall, λ determines the range over which the probability curve for the second cell is nonzero.

**Figure 4:**
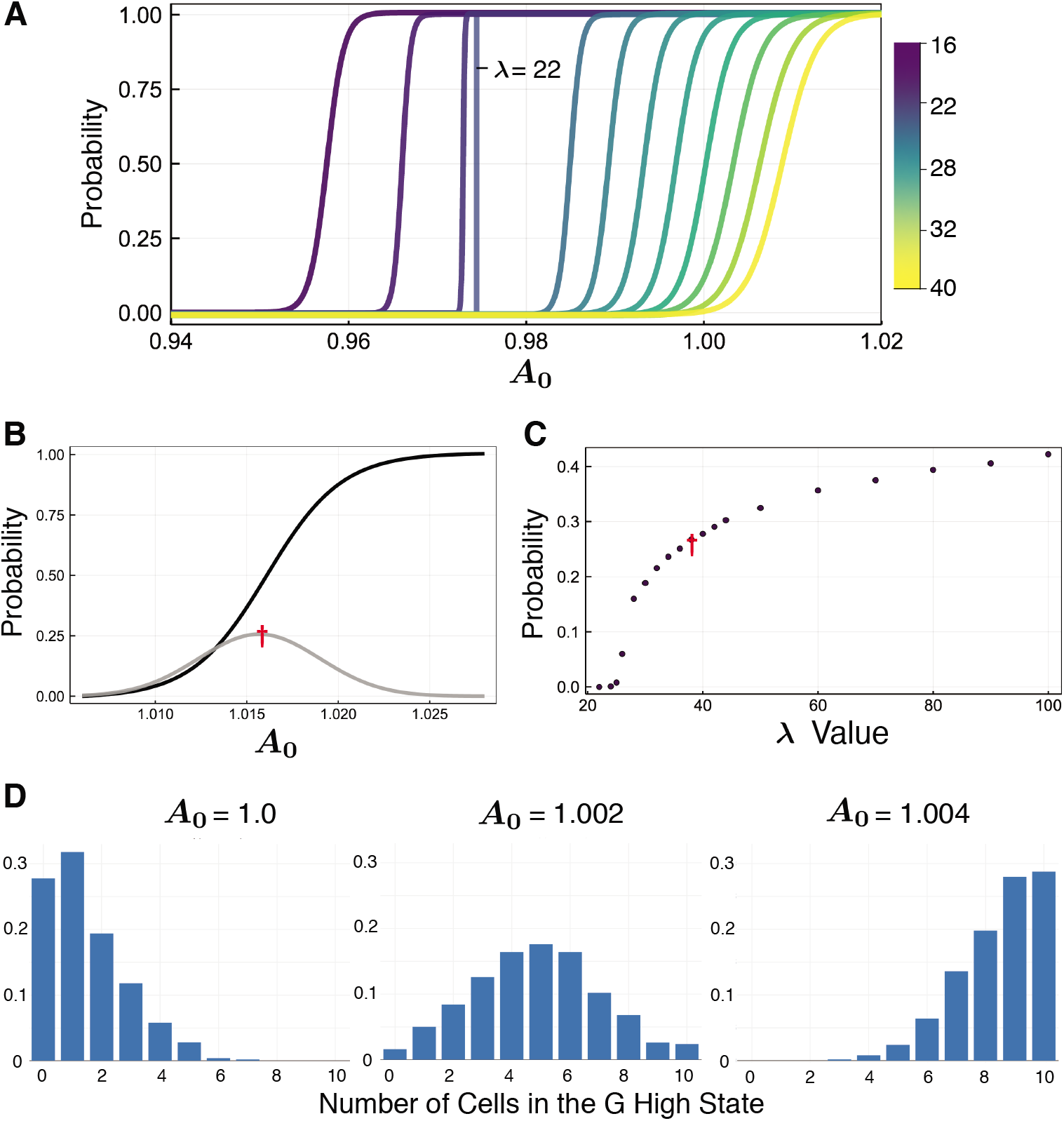
**(A)** Probability distributions of cell fate commitment (to *G* high state) for cell 2 in a chain of two cells for λ ∈ [16,40]. **(B)** Probability distributions for a loop of two cells as a function of *A*_0_, with λ = 38. Probability of cell fate commitment to *G* high state (black); probability that the two cells will not commit to the same fate (gray), with the maximum of the distribution marked (red cross). **(C)** For a loop of two cells, the maximum probability that the cells do not converge to the same lineage as λ varies; red cross equivalent to (B) is marked for comparison. **(D)** Comparison of distributions for the number of cells that commit to the *G* high state in a loop of 10 cells with λ = 30.

We next assessed how different values of λ changed coordination of cell fate decisions between lineages. Previously, we observed that for any trajectory of a loop topology of any size, where λ = 1, all cells converged to the same state, regardless of the value of *A*_0_. Further, although all cells in the topology converge to the same lineage, the probability that they all converge to a given lineage depended on *A*_0_. To see whether or not this behavior was conserved for other values of λ, we tested a two cell loop topology with a range of λ values. We observed that there exists a value λ* ≈ 22 such that for λ < λ*, all the cells coordinate their lineage just as we had seen before. Interestingly, for λ > λ*, the two cells do not always coordinate their lineage decisions. For example, Fig. S6 gives sample trajectories showing two cells in a loop converging to each combination of states. Fig. 4D shows the two cell loop results for λ = 38, giving the probabilities that each cell converges to the *G* high state as well as the probabilities that the cells are decoupled, converging to opposite lineages. For each λ, we recorded the maximum value of the probability that the cells converge to different lineages. Fig. 4C shows how the maximum probability of cells converging to different lineages increases with λ.

Next, we looked at larger loops of 10 cells. For each value of *A*_0_, we simulated 500 trajectories and recorded how many cells were in the *G* high state verses the *G* low state. Fig. 4D shows results for λ = 30 and *A*_0_ = 1.0, 1.002, and 1.004. We see that the distribution of cells converging to each state changes with the value of *A*_0_. Further analysis of distributions with different values of λ can be found in Fig. S7.

We have identified an explicit mechanism by which cell-cell communication can break the symmetry of a homogeneous population of progenitor cells, and give rise to stable, heterogeneous populations of lineage-committed cells. The proportion of cells committed to each lineage depends on the external environment, *A*_0_, and the strength of cell-cell coupling due to signaling, λ. Moreover, these results show that fluctuations in the external environment can lead to shifts in the relative abundances of committed cell types. These results corroborate previous work that studied the generation of heterogeneous cell populations through stem cell differentiation [34], and showed that external signals (e.g. through cell-cell communication) are required both to maintain heterogeneous cell populations and to shift relative cell types abundances in response to environmental perturbations.

### Intrinsic and extrinsic noise alter cell fate decision-making boundaries

We have until now assumed that signals are passed between cells with perfect fidelity. In fact, multiple sources of noise contribute to imperfect communication between cells, and the modeling framework here lends itself well to the investigation of the effects of intrinsic vs. extrinsic noise [35, 36]. We investigated the impact of two different sources of noise in cell-cell communication: due to cell-extrinsic factors, i.e. noise with respect to the extracellular environment (Eq. 6); or due to cell-intrinsic factors, i.e. noise due to signal transduction downstream of a paracrine signaling factor (Eq. 7). These sources of noise are represented in the model by varying either the baseline level of cell-cell communication (extrinsic noise) or the cell signaling pulse level, i.e. the intrinsic signal transduction noise (Fig. 5A).

**Figure 5:**
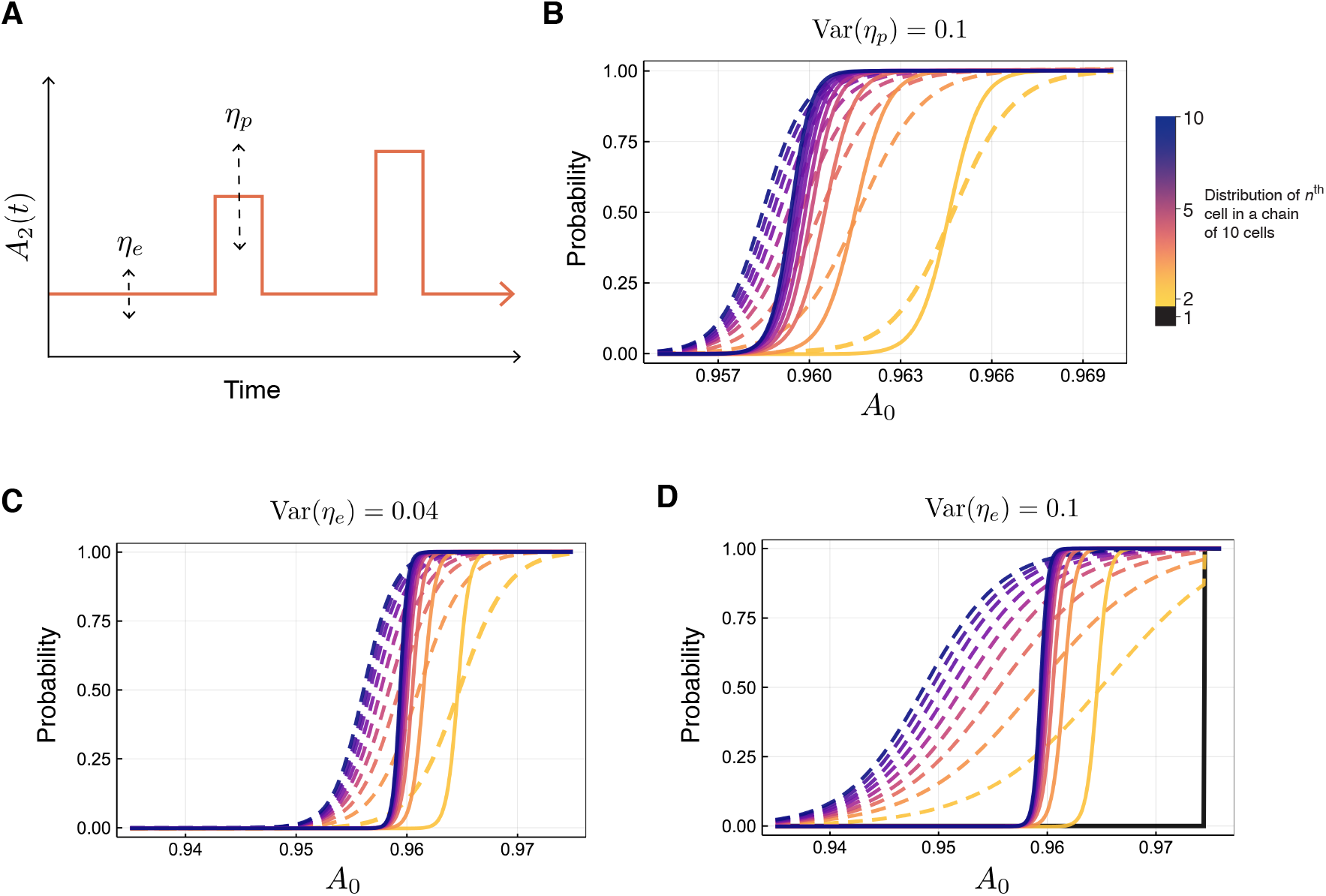
**(A)** Schematic depicting how extrinsic noise (*η_e_*) and intrinsic noise (*η_p_*) are modeled through their effects on the signaling strength *A*. **(B)** Probability distributions of cell fate commitment (to *G* high state) for each cell in a ten cell chain, with λ = 18: no added noise (solid lines); and with added noise (dashed lines). The colors darken as the position of the cell along the chain increases. The type of noise modeled is intrinsic, with 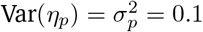. **(C)** As for (B), but for extrinsic noise modeled, with 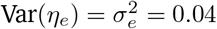. **(D)** As for (B), but for extrinsic noise modeled, with 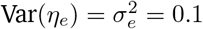.

For simple topologies, as we would expect, the observed variability in cell fate outcomes increases as the variance of either the extrinsic or the intrinsic noise increases (Fig. S8). This results also holds for larger cell-cell communication topologies, e.g. a ten-cell loop, intrinsic (*η_p_*) and extrinsic (*η_e_*) noise (Fig. 5B-D) both reduce the sensitivity of the cell fate decision-making boundary. We also observe unexpected and striking results. First, not only are the probability distributions flattened by either noise source, but they are shifted to the left, i.e. intrinsic and extrinsic noise directly affect the decision-making boundary (by shifting its mean), as well as the sensitivity of cell-fate decision-making. Second, even under the presence of noise, we still observe the effects of cooperativity at play through the sharpening of distributions down a chain of cells (Fig. 5D). That is, individual noisy signals increase the variability (reduce the sensitivity) of cell fate decision-making, but the cumulative effects of noisy signaling can at least partially compensate for this, and decrease the variability in cell fate decision-making.

Through comparison of the relative effects of the intrinsic signal transduction noise (Fig. 5B) and the extrinsic extracellular noise (Fig. 5C-D), we see that the impact on the cell fate decision-making boundary is much larger for extrinsic rather than intrinsic noise contributions. From inspection of Eqs. 6–7, this is in part due to smaller duration of the pulse *τ* relative to the mean wait time *μ*. The result of which is that when 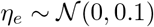, (Fig. 5D), the probability curves for cells 2-10 in the chain flatten to the extent that the sensitivity with which to distinguish cell fate decision-making by cell position along the chain is lost entirely. Moreover, the probability curves for cells ≥ 2 along the chain intersect with the point at which cell one (which is deterministic) switches fates from the low to the high state (black line in Fig. 5D), thus forcing all other cells also into the high state with probability = 1. This coordination of cell fates is influenced by cell-cell coupling (here we use λ = 18 < λ*, i.e. a value of λ for which we observe complete coupling (fate coordination) between cells). The dominant impact of extrinsic over intrinsic noise here is in agreement with previous works, including a study that quantified the contributions of extrinsic and intrinsic noise in the MAPK signaling pathway, and showed that extrinsic noise is the dominant driver of cell-to-cell variability [37]. It has also been shown that explicit extrinsic noise contributions are necessary to explain mRNA abundance distributions [38].

In summary, the effects of intrinsic and extrinsic noise on cell-cell communication topologies are to increase the variance of the resulting cell fate distributions and (surprisingly) to alter the mean values of these probability distributions. In other words, the presence of noise alone can force cells to change lineages. The observed increases in the variability of cell fate decision-making are maintained for large cell-cell communication topologies. A similar result was described in a study of cell fate decision-making during early mouse gastrulation [39]: transcriptional noise is greatest at the point of cell fate decision-making (when epiblasts begin to differentiate). Our results reiterate the same point made by Mohammed et al., that gene expression noise is beneficial during windows of cell fate decision-making as it leads to an increased possible repertoire of cell fates. Our findings go further in that they suggest a rationale: that the increase in transcriptional noise results from noisy extracellular factors influencing cell-cell signaling during differentiation.

## Discussion

Despite many theoretical and experimental advances in our understanding of gene regulatory network (GRN) dynamics, our ability to use GRN models to explain cell fate decision-making during differentiation of multipotent progenitor cells remains limited. Here, with application to a well-studied cell fate GRN – the *GATA1-PU.1* mutual inhibition loop [10–14] – we introduced a new model that can simultaneously describe GRN dynamics and single cell-resolved cell-cell communication. Notably, although cell-cell communication is often assumed to be a critical component of cell differentiation, it is rarely incorporated into models. The previous studies that have characterized cell-cell communication in models did not capture the detailed complexity of these dynamics, by making simplifying assumptions either regarding the GRN dynamics [23] or regarding the mechanisms by which cells signal [21, 22]. We found that by combining these dynamic processes that describe noisy cell fate decision-making at single-cell resolution we were able to reconcile several outstanding controversies in the field.

Over a large domain of possible cell-cell communication topologies, we found that cell-cell communication alters cell fate decision-making boundaries, which become probabilistic in response to the levels of external signaling factors. This helped to reconcile a controversy: that of whether or not transcriptional stochasticity is sufficient to initiate the granulocyte-monocyte vs. megakaryocyte-erythrocyte cell fate decision. Previous models supported the hypothesis, however Hoppe et al. presented compelling evidence to contradict it [15]. Our results agree with Hoppe et al., in that we show that eventual cell fate cannot be inferred from the initial gene expression state alone—and go further in emphasizing the stochastic nature of these cell fate decisions [40]. Analysis of cell-cell coupling led to the discovery that stable distributions of heterogeneous cell types can be robustly maintained by a set of external signals. This result offered insight into another open question: that of how population-level cell fate behaviors emerge during cell differentiation [34]. Finally, we showed how (primarily extrinsic) noise increases the variability of cell fate decision-making, in line with previous analyses of transcriptional noise during development [39].

We sought to tightly constrain model complexity here, for reasons of parsimony and interpretability. Relaxing some of the constraints imposed will likely also yield interesting results. We assumed throughout that cells were initially homogeneous, i.e. they shared the same initial conditions and internal GRN networks. Heterogeneous initial conditions and heterogeneous cell fate decision-making (i.e. different GRNs in different cells) ought to be explored, for example by considering interactions between two progenitor cell types, each controlled by their own GRN. There is also much room for exploration of larger and more varied cell signaling network topologies. Here, future work should be guided by data, as it becomes harder to justify large signaling networks chosen a priori. For example, spatial transcriptomics [41] could be used to infer cellular networks (e.g. of cells of similar type) that could then be input to our model framework. There ought to include dissensus as well as consensus signals. There are also a variety of well-informed modifications that could be made to the signaling model definition. For example, currently signals only interact with *GATA1*; it would be interesting in future work to explore signals that also interact directly with *PU.1* expression by modifying the model parameter *B*.

A central challenge for the model introduced here is that of fitting to data. Ideally this would require both spatially and temporally resolved single-cell transcriptomic data—at the limits of current technologies, although this is changing [42]. Thus in the current work we rely on comparison of qualitative features arising from the model with previous experimental studies. Moreover, due to the hybrid deterministic-stochastic formulation of the model, resulting in time-dependent signaling parameters, we doubt that it will be possible to derive an appropriate likelihood function for Bayesian parameter inference. Thus, approximate Bayesian computation will likely be required. Yet, even here, simulation times may be prohibitive or require further approximations to be made.

Future work should also treat more carefully the stochastic nature of gene expression, for example by replacing the deterministic GRN dynamics with discrete stochastic simulation or stochastic differential equations (SDEs). This would be straightforward to accomplish numerically, although it will complicate the analysis of model bifurcations. While the complete tractability of the deterministic model made it an ideal first candidate with which to study the effects of cell-cell communication, noise undoubtedly plays important roles in regulating single cell phenotypes [43].

In conclusion, the introduction of a single-cell resolved cell-cell communication model of GRN dynamics has helped to explain numerous cell fate decision-making phenomena. Even in a tightly constrained model space, we have shown that changes in the distribution of cell fates due to cell-cell communication can be broad and varied. More generally, we have highlighted the need to consider multiscale effects in models of cell fate dynamics. We anticipate that the application of these methods to different GRNs will lead to advances in our understanding of specific cell fate decision-making control points, as well as general principles describing the control of stem cell differentiation.

## Acknowledgements

We would like to thank Rong Lu and members of the Lu lab for many helpful discussions. We are grateful to Michael P.H. Stumpf for insightful comments and feedback on the manuscript. This work was partially supported by a USC WiSE Top-up Fellowship (to M.K.F.) and the National Science Foundation (DMS 2045327 to A.L.M.).

## Software and data availability

The model was developed in Julia, and all the code associated with simulation and analysis is available under an MIT license at: https://github.com/maclean-lab/Cell-Cell-Communication.

