## Supplementary Figures for "A single-cell resolved cell-cell communication model explains lineage commitment in hematopoiesis"

### Supplementary Information

| Parameter | Description | Value |
|---|---|---|
| $\alpha_1$ | Activation of $G$ by $A$ | 1.0 |
| $\alpha_2$ | Activation of $G$ by self | 0.25 |
| $\alpha_3$ | Activation of $P$ by $B$ | 1.0 |
| $\alpha_4$ | Activation of $P$ by self | 0.25 |
| $\alpha_5$ | Activation of $X$ by $G$ | 0.01 |
| $\beta_1$ | Degradation of $G$ | 0.01 |
| $\beta_2$ | Degradation of $P$ | 0.01 |
| $\beta_3$ | Degradation of $X$ | 0.01 |
| $\gamma_1$ | Inhibition of $G$ by $A$ | 1.0 |
| $\gamma_2$ | Inhibition of $G$ by self | 0.25 |
| $\gamma_3$ | Inhibition of $G$ by complex $GP$ | 1.0 |
| $\gamma_4$ | Inhibition of $P$ by $B$ | 1.0 |
| $\gamma_5$ | Inhibition of $P$ by self | 0.25 |
| $\gamma_6$ | Inhibition of $P$ by complex $GP$ | 1.0 |
| $\gamma_7$ | Inhibition of $P$ by complex $GX$ | 1.0 |
| $\gamma_8$ | Inhibition of $X$ by $G$ | 0.01 |
| $\gamma_9$ | Inhibition of $X$ by $C$ | 10.0 |
| $B$ | External signal to $P$ | 0.5 |
| $C$ | External signal to $X$ | 0 |

Table S1: Parameter values used for the simulation of GRN dynamics. Source: Chickarmane et al. [1]

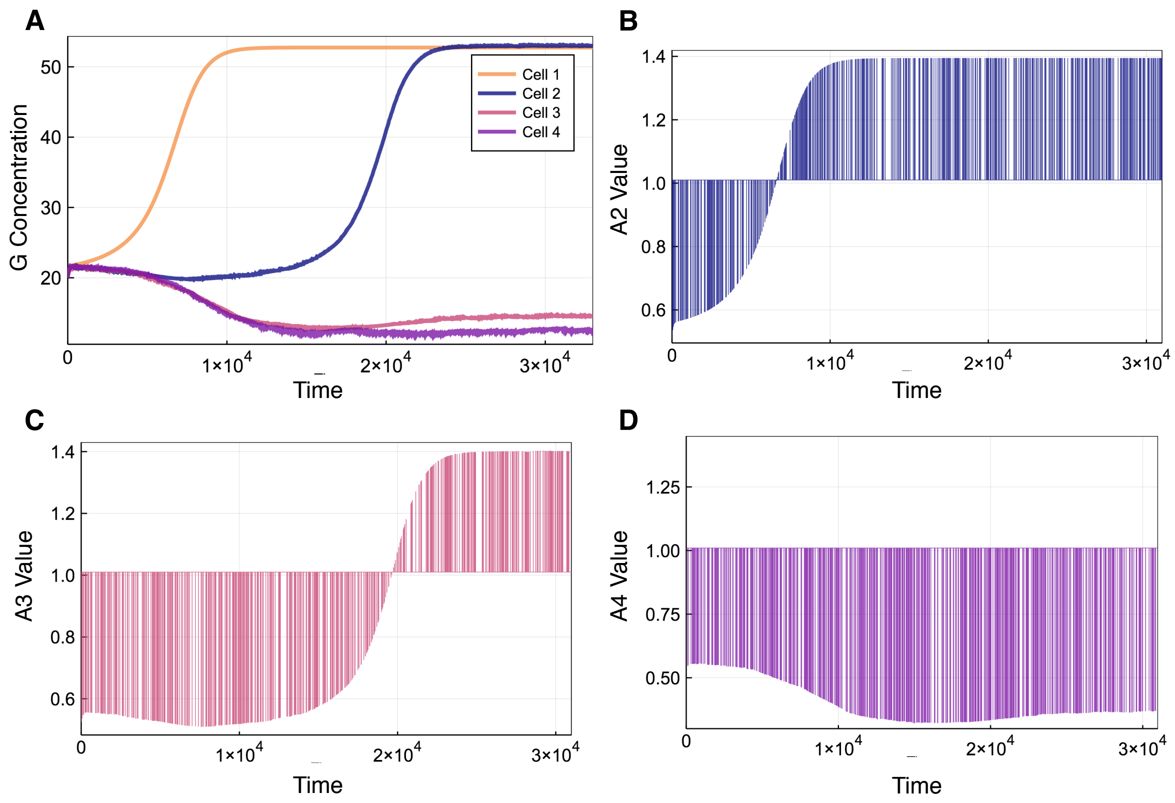


Figure S1: (A) Sample trajectory of a chain of four cells where $A_0 = 1.01$ and $\lambda = 38.0$. (B)-(D) Plots of $A_2(t), A_3(t)$, and $A_4(t)$ over the same timescale as the trajectory.

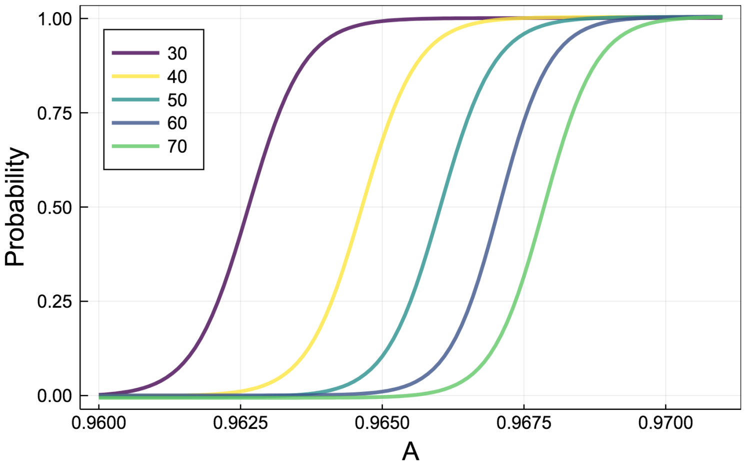


Figure S2: Probability distributions of cell 2 in a chain of cells converging to the $G$ high state where $\lambda = 18$ with different mean wait times, $\mu$, given in the legend.

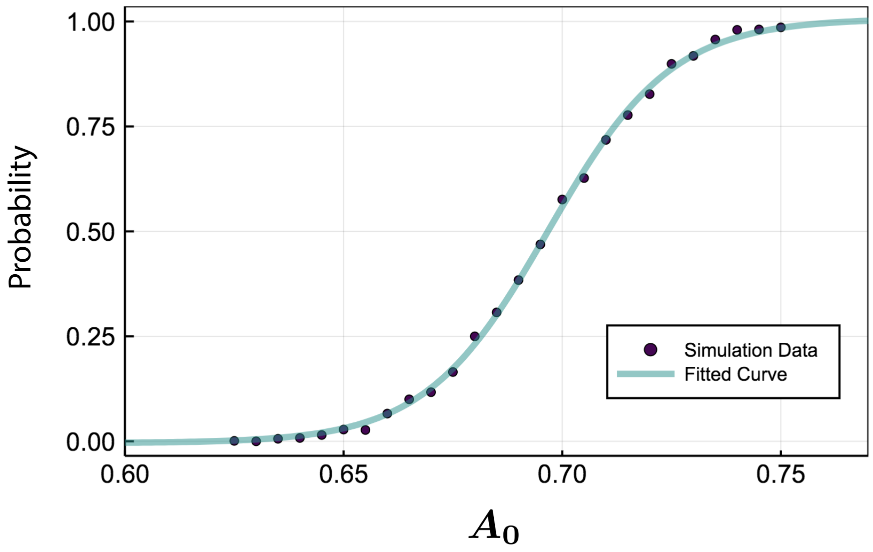


Figure S3: Sample of simulated data points along with the fitted curve for cell 2 in a chain of cells where $\lambda = 1$.

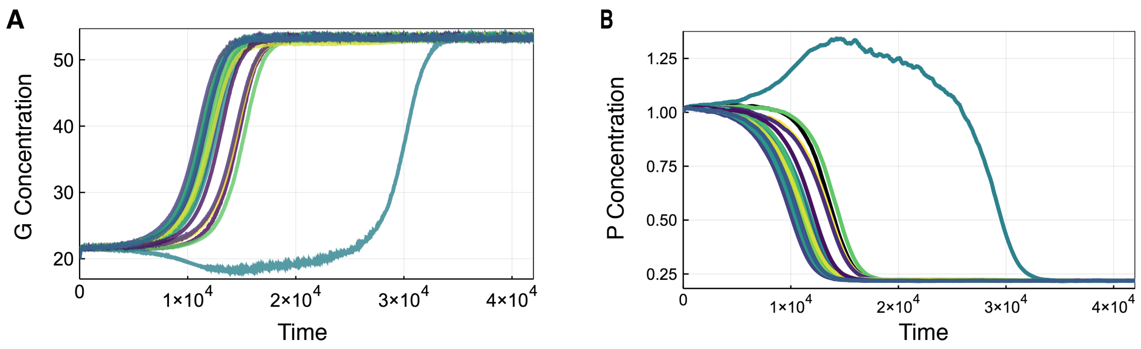


Figure S4: Sample simulation of a loop of 20 cells with concentrations of both $G$ and $P$ over time, where $\lambda = 28.0$ and $A_0 = 1.0$.

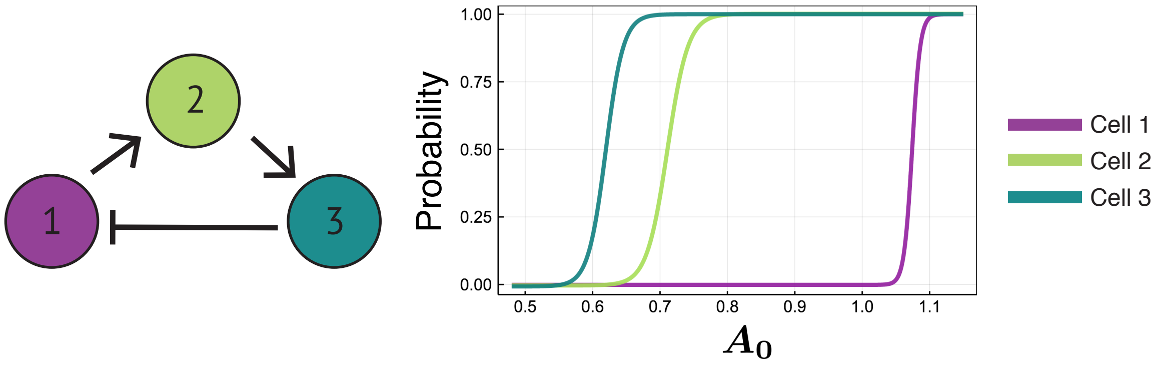


Figure S5: Cell fate probability distributions for a loop of three cells with one inverse signal.

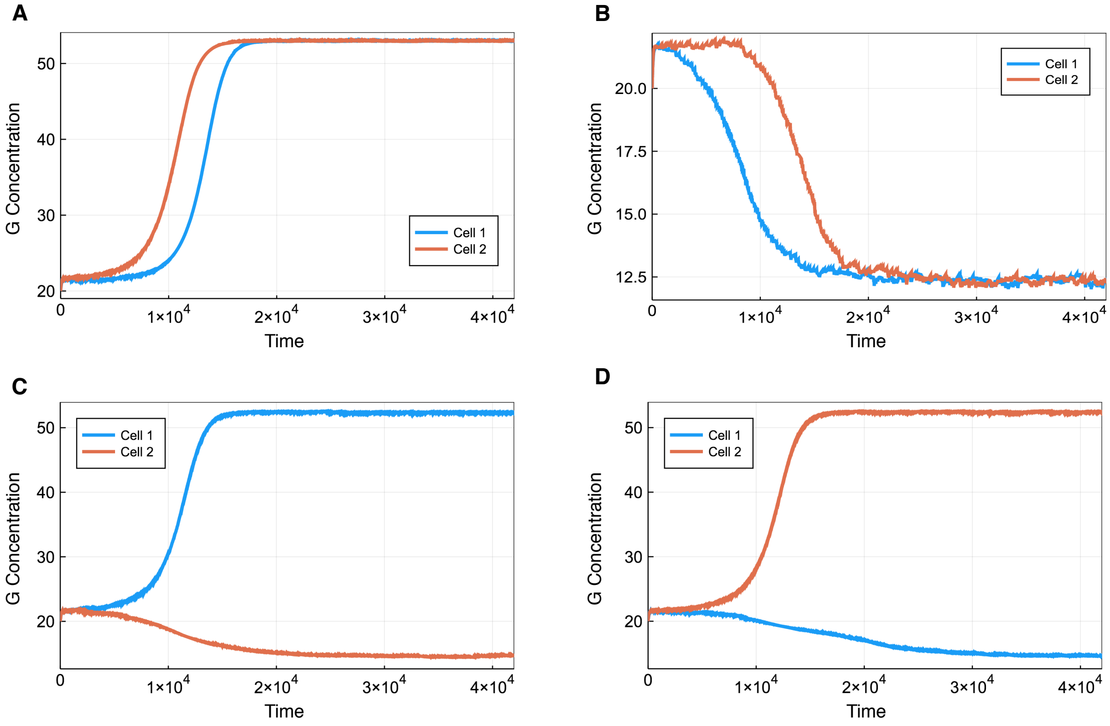


Figure S6: (A)-(D) Sample trajectories of a loop of two cells with $A_0 = 1.0175$ and $\lambda = 40$.

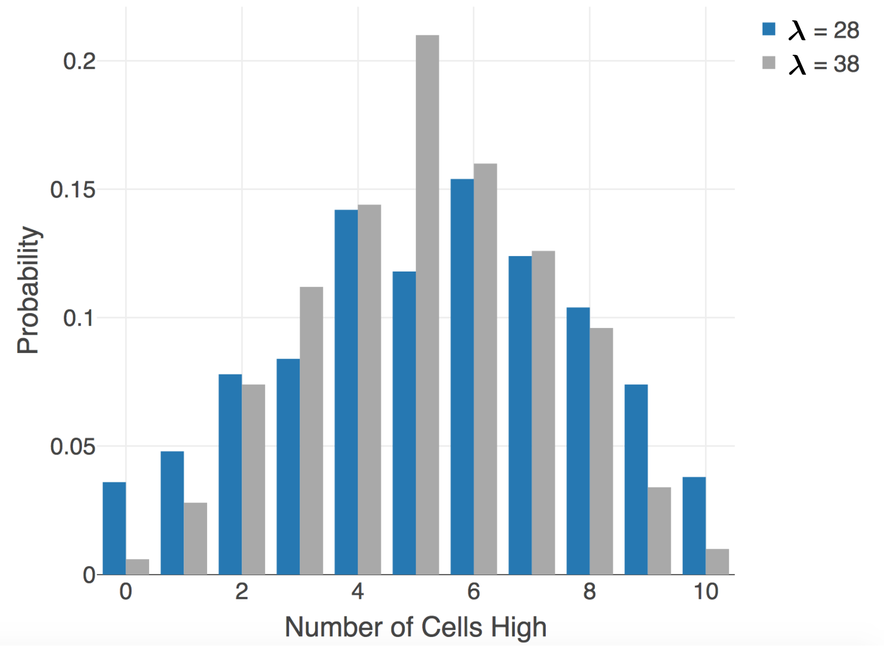


Figure S7: Distributions of loops of 10 cells with different values of $\lambda$. The values of $A_0$ were selected so that the distributions had similar expected values. For the blue distribution, $A_0 = 0.9973$ and the expected value is 5.284. For the grey distribution, $A_0 = 1.016$ and the expected value is 5.154

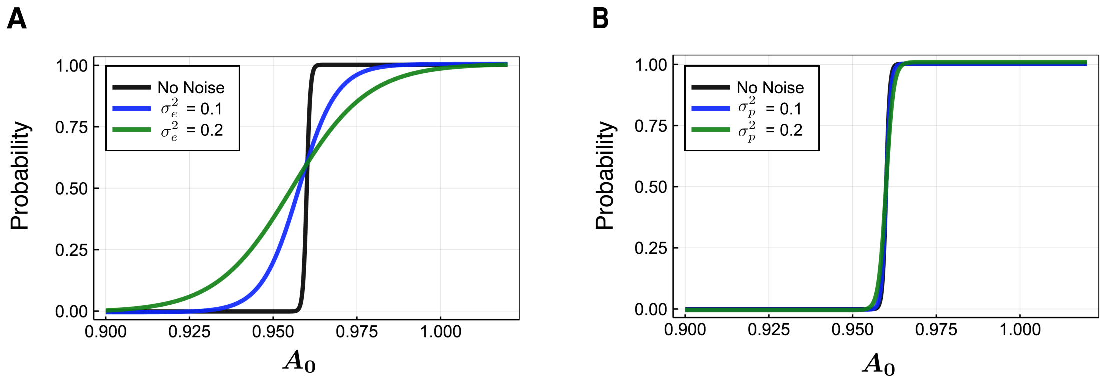


Figure S8: Probability distributions that a given cell in a two cell loop with $\lambda = 18$ will converge to the erythroid state with varying amounts of (A) extrinsic noise or (B) intrinsic noise.
